# Estimating steady-state evoked potentials in the limit of short data duration and low stimulation frequency

**DOI:** 10.1101/2024.05.26.595940

**Authors:** Marco Buiatti, Davide Saretta

## Abstract

Because of their high signal-to-noise ratio and robustness to artifacts, Steady-State Evoked Potentials (SSEP) - the periodic responses elicited by periodic stimulation designs - are increasingly used in human neuroscience for measuring stimulus-specific brain responses in a short presentation time. While widely applied to measure sensory responses with stimulation frequencies higher than 8 Hz, they are also successful to investigate high-order processes and/or early development characterized by slower time scales, requiring very low stimulation frequencies around 1 Hz. However, applications of these low frequency paradigms on developmental or clinical populations, typically relying on very short data recordings, pose a methodological challenge for SSEP estimation. Here we tackled this challenge by investigating the method of analysis that most efficiently compute SSEP at low stimulation frequencies in the limit of short data, and by estimating the minimum data length necessary to obtain a reliable response. We compared the performance of the three most commonly used measures of SSEP (power spectrum (PS), evoked power spectrum (EPS) and inter-trial coherence (ITC)) at progressively shorter data segments both on simulated data and on EEG responses to on-off checkerboard stimulation at two ‘low’ frequencies (4 Hz and 0.8 Hz). Results, consistent between simulated and real data, show that while for long data length EPS and ITC outperform PS, for short data length the three measures are equivalent, and the crucial parameter is the length of the sliding window over which each measure is computed: the longer the better for PS and EPS, whereas the opposite occurs for ITC. For the analysed dataset, the shortest data length required to estimate a reliable SSEP is as short as 8 cycles of stimulation, independently from the stimulation frequency. This study provides practical indications for reliable and efficient application of low-frequency SSEP designs on short data recordings.

## 1 INTRODUCTION

Electroencephalography (EEG) and magnetoencephalography (MEG) critically contribute to our knowledge about sensory, perceptual and cognitive brain functions by providing direct, non-invasive measures of brain responses to exogenous stimuli. However, obtaining stimulus-related brain responses from EEG and MEG (hereafter MEEG) recordings is challenging because of the interference of ongoing, stimulus-unrelated MEEG activity and of artifacts of biological and technical origin. One experimental approach that minimizes these sources of interference consists in presenting the stimuli in strictly periodic sequences at a specific (“tag”) temporal frequency and exploits the property of the brain to respond periodically at the same tag frequency (Norcia et al., 2015; Picton et al., 2003). These periodic responses are traditionally named “steady-state evoked potentials” (SSEP) because of the stability of their amplitude and phase in response to periodic stimulation (Regan, 1966). While some researchers consider this definition appropriate only for stimulation frequencies sufficiently high to elicit a nearly sinusoidal response, we adopt the wider meaning proposed by (Norcia et al., 2015) for which responses are called SSEP if both the stimulus and the response are periodic, a definition that includes the low frequency stimulations studied in this work. These experimental designs have also been named “frequency-tagging” when using simultaneous multiple stimulation (“tag”) frequencies to tag distinct features of the stimuli within the same presentation (Buiatti et al., 2009; Tononi et al., 1998).

SSEP are typically manifested in the MEEG recordings by a sharp peak in the signal’s power spectrum at that specific frequency. Since the ongoing MEEG activity is broadband in frequency, the stimulus-related response in the frequency domain is easily discriminated from the stimulus-unrelated activity, yielding a much higher Signal-to-Noise Ratio (SNR) than the one obtained in the time domain with the repetitive presentation of single, temporally isolated stimuli (event-related potentials), because the time scales of stimulus-related responses overlap with those of the ongoing neural activity. Moreover, since most MEEG artifacts (eye movements, blinks) are also broadband in frequency, SSEP are more robust to artifacts and require a lighter artifact rejection procedure than event-related potentials (Kumaravel, Farella, et al., 2022).

SSEP are widely used to investigate sensory processing, especially in the visual domain (Steady-State Visually Evoked Potentials—SSVEP (Norcia et al., 2015)) and in the auditory domain (Auditory Steady-State Responses—ASSR (Picton et al., 2003)). Typical presentation rates depend on the frequency ranges in which these sensory systems are most responsive: 8–20 Hz for SSVEP and around 40 Hz for ASSR. In these frequency ranges, it is possible to obtain very high SNR, both because due to the intrinsic temporal properties of the underlying neural circuits the brain response is almost sinusoidal and its amplitude is high (Norcia et al., 2015), and because the amplitude of both ongoing EEG activity and biological EEG artifacts is relatively low.

Recent studies have extended the use of SSEP to lower frequencies (0.5–6 Hz, encompassing the delta and theta frequency ranges) either because they focused on higher-level neural processing characterized by longer temporal scales (e.g., syllables and words in speech perception (Buiatti et al., 2009; Ding et al., 2016; Kabdebon et al., 2015), or because the experimental design was based on the infrequent presentation of key stimuli among control ones (e.g., selectivity for faces among other visual objects (de Heering & Rossion, 2015; Rossion et al., 2015), or to cope with the slow neural processing due to the immaturity of the visual system (e.g., face perception in newborns (Buiatti et al., 2019)). Obtaining reliable brain responses within this low-frequency range is more challenging for at least three reasons: 1) Low-frequency SSEP consist in a sequence of transient responses rather than continuous sinusoidal ones usually resulting in significantly lower spectral amplitudes (Norcia et al., 2015; Picton et al., 2003); 2) The interference of both EEG ongoing activity and artifacts increases for decreasing frequencies; 3) Since oscillation cycles are longer, we need lengthier data to capture the related oscillatory responses.

The latter requirement is particularly problematic in studies involving subjects with limited attention, especially in developmental studies using visual stimulation where the attentional span that may not exceed a few tenths of seconds (Buiatti et al., 2019).

The purpose of this work is to identify the method of analysis and the parameters that most efficiently compute SSEP at low stimulation frequencies in the limit of short data, and to provide an estimate of the minimum data length necessary to obtain a reliable response for a given stimulation frequency.

For this purpose, we tested the influence of data length for low stimulation frequencies on the three most common spectral methods of SSEP estimation:

1. The Power Spectrum (PS): It measures the amplitude of the oscillations of the recorded signal at the stimulation frequency; in the low frequency range, in order to isolate the frequency peak from the ongoing stimulus-unrelated activity, the PS is typically normalized by dividing the amplitude at the stimulation frequency by a fit of the background activity at neighbouring frequency bins (Picton et al., 2003);
2. The Evoked Power spectrum (EPS): Assuming that SSEP are time-locked to the periodic stimulation, the EEG is first time-averaged to cancel out the non-time-locked brain activity; the power spectrum is then computed on this evoked response, and normalized as the PS (Ding et al., 2016; Rossion et al., 2015);
3. The inter-trial coherence (ITC): Assuming that SSEP are phase-locked to the stimulation, the ITC computes the coherence of this phase-locking along the stimulation, independently from the amplitude of the oscillations at the stimulation frequency (Forget et al., 2010; Kabdebon et al., 2015). Although ITC is already a normalized measure (Tallon-Baudry et al., 1996), some researchers apply a frequency normalization to isolate a frequency-specific effect (Fló et al., 2022).

All the three methods share the first key step: choosing the length *wl* of the sliding time window on which to compute the measure. In order to have a frequency bin corresponding to the stimulation frequency, *wl* must be a multiple of the oscillation cycle (Bach & Meigen, 1999). Given a finite-size EEG signal, the choice of *wl* is crucial because it sets the tradeoff between the number of samples (higher for shorter *wl*) and the frequency resolution (higher for longer *wl*). This choice is even more crucial for data vulnerable to artifacts like developmental or clinical recordings, where choosing too long *wl* might imply a drastically high rejection rate.

We therefore tested the influence of decreasing data length on the three measures by varying two parameters of crucial importance: the *wl* and the normalization method.

To explore the general behavior of the three measures in the short data limit, we first tested them on data simulated with a generative model matching the basic features of real SSEP and background EEG activity. We then validated the results on a real EEG dataset including SSVEP recorded from the presentation of an on-off checkerboard at two stimulation frequencies: 4 Hz, at the lower border of the frequency range for which the single-trial response is an almost continuous oscillation following the periodic stimulation, and 0.8 Hz, characterized by transient responses and wide-amplitude ongoing fluctuations.

Based on these results, we provide a set of methodological indications to efficiently compute SSEP in the low frequency range and short data limit.

## 2 MATERIALS AND METHODS

### 2.1 EEG dataset

Twelve right-handed (Edinburgh Inventory) native Italian speakers participated in the experiment (5 females; 21–29 years, mean age 25 years). All participants had normal or corrected-to-normal visual acuity, and reported no history of neurological or psychiatric disorders. All participants provided informed written consent to take part in the experiment, which was approved by the Ethical Committee of the University of Trento (Italy). The dataset was recorded as a “toy” dataset to develop data analysis tools (Montagna et al., 2017) for visual frequency-tagging paradigms to be used in newborn/infant EEG studies (Buiatti et al., 2019).

Stimuli consisted of a black and white 10x10 square checkerboard subtending approximately 15° by 15° of visual angle on a uniform grey background, presented at a distance of 90 cm from the subject’s eyes. Stimuli were presented with a sinusoidal on–off 100% contrast temporal modulation (the visibility of the black/white checkerboard gradually rises with respect to the gray background from 0% at the beginning of the cycle to 100% at mid-cycle, then gradually decreases to 0% towards the end of the cycle) at 2 frequency rates (0.8 Hz and 4 Hz) in two blocks of 40 s, using the Psychtoolbox 3.0.12 for Windows in Matlab (The MathWorks Inc., Natick, Massachusetts). We used sinusoidal contrast modulation instead of squared on–off dynamics both to minimize nonlinear effects in the brain frequency response (Norcia et al., 2015) and because it is a more pleasant and less fatiguing visual stimulation than a squarewave stimulation mode for the subjects (Buiatti et al., 2019). Subjects were asked to fixate at the center of a grey diagonal cross overlapped to the checkerboard. Subjects were presented with additional stimulation protocols involving face and number perception; the associated data were not analysed in the present study since they are not relevant for its purpose.

The experiment was performed in an electrically shielded and sound-attenuated cabin. EEG was recorded with a BrainAmp amplifier (Brain Products, Munich) using 64 Ag/AgCl sintered ring electrodes mounted in an elastic cap (Easycap, Munich) and placed equidistantly according to the 10/20 system, with a vertex reference (Cz) and ground electrode in AFz. Electrode impedances were kept below 15 kΩ. Data were sampled at 500 Hz and analog filtered between 0.016 and 250 Hz during recording.

#### EEG data preprocessing

Continuous raw data were imported in the EEGLAB software (Delorme & Makeig, 2004) and band-pass filtered between 0.1 and 40 Hz with the default EEGLAB filter to remove DC and high-frequency noise. Data were segmented in windows corresponding to the stimulation blocks. To avoid the interference of the first transient response to visual stimulation at the onset of each block and to allow entrainment to take place, we removed the first 1.25 s (corresponding to one stimulation cycle) from the 0.8 Hz blocks and the first 0.5 s (corresponding to two stimulation cycles) from the 4 Hz blocks before further analysis. Each segment was visually inspected and portions corresponding to integer stimulation cycles containing nonstereotyped paroxysmal artifacts were discarded. Bad channels were identified with the LOF algorithm (Kumaravel, Buiatti, et al., 2022; Kumaravel, Farella, et al., 2022) and discarded (no more than two per subject). To identify and remove stereotypical artifacts, the default EEGLAB ICA decomposition was computed on the concatenation of all segments. Blinks, eye-movements and other topographically localized artifacts were discarded by removing the corresponding independent components identified by ADJUST, an algorithm for automatic detection of artifacted ICA components (Mognon et al., 2011). Further artifacts were discarded by removing related ICA components identified by visual inspection of their topography and spectro-temporal profile. EEG signals in bad channels were interpolated with the EEG signals from neighbouring channels (standard spherical interpolation method in EEGLAB), and the resulting clean data were re-referenced to average reference.

For each subject and each stimulation frequency, the first pre-processed data block was selected for further analysis. Since the longest duration of the pre-processed data blocks compatible with all subjects was 25 s (a data length widely sufficient for the purpose of this work), we did all the analyses on 25 s data segments for each subject and stimulation frequency.

### 2.2 Simulated data

We generated the simulated data by summing two components:

*Background component:* Stimulus-unrelated ongoing EEG activity was generated as 1/f-like noise (power spectrum increasing as 1/*f*^*α*^ for *f* → 0), where the power-law exponent *α* is estimated from the resting state segments of the EEG dataset as follows: 1) for each subject and electrode, we computed power spectrum of the EEG signal between 0.5 Hz and 4.5 Hz; 2) for each subject, we estimated the power-law exponent from the linear fit in log-log scale of the average of the power spectrum over the posterior cluster of electrodes *pROI* indicated in Section 2.4). The resulting power-law exponent was *α* = 1.4 ± 0.24, fully compatible with the literature (Buiatti, 2023). Therefore, we generate 1/f-like noise with power-law exponent *α* = 1.4 by using the AR model proposed by (Kasdin, 1995) (Eq. 116), as implemented in the Matlab function *dsp*.*colorednoise* (The MathWorks Inc., Natick, Massachusetts).

*SSEP component:* We simulated an SSEP for each of the two stimulation frequencies by a combination of single-trial transient evoked responses (Capilla et al., 2011) generated as Gaussian “peaks”, each one determined by its parameters of latency, width and amplitude (Krol et al., 2018). For each stimulation frequency, we estimated the parameters by modeling the transient responses as the two most prominent waves in the Event-Related Potentials (ERP) time-locked to the beginning of each stimulation cycle, computed from the real EEG data, averaged over the posterior cluster of electrodes *pROI* indicated in Section 2.4 and over all subjects. For the 0.8 Hz stimulation frequency, we generated a negative wave at 200 ms (width: 50 ms) and a positive wave at 450 ms (width: 100 ms). For the 4 Hz stimulation frequency, we generated a negative wave at 60 ms (width: 30 ms) and a positive wave at 200 ms (width: 50 ms). To simulate single-trial variability, for each trial all parameters (latency, width and amplitude) varied according to a Gaussian with a width of 20% with respect to the set value (Krol et al., 2018).

For each stimulation frequency, we generated simulated data with three values of signal-to-noise ratio (SNR), obtained by varying the ratio between the background component and the SSEP component. We set the “middle” SNR value such that the normalized power at the stimulation frequency (NP, see Section 2.3.1) matched the one measured on real, artifact-free EEG data, averaged over *pROI*, the posterior cluster of electrodes selected for the analysis (Section 2.4) and over all subjects, for segments of 20 cycles of oscillation (f=0.8 Hz, 25 s data length: SNR=2.8; f=4 Hz, 5 s data length: SNR=3.1). We set the “low” and “high” SNR value by respectively dividing and multiplying the ratio between the background component and the SSEP component corresponding to the middle SNR by the factor 1.4.

Hereafter we refer to the simulated data as “simulated response”, and to the background component only as “simulated rest”.

### 2.3 SSEP measures

We tested the three most popular spectral measures of SSEP: the power spectrum (PS), the evoked power spectrum (EPS) and the intertrial coherence (ITC). For both the real EEG data and the simulations, the signal consists of a single continuous segment of data with variable length for each subject.

#### 2.3.1 Power Spectrum

To compute the PS, the signal was first segmented in partially overlapping epochs (overlap= 50%) obtained by sliding a window of length *wl* along the data. The Fourier transform *F*(*f*) of each epoch was then calculated using a fast Fourier transform algorithm (MATLAB function FFT). For data shorter than *wl*, zero-padding to *wl* was applied before FFT. The power spectrum was calculated from these Fourier coefficients as the average over all epochs of the single-epoch power spectrum:

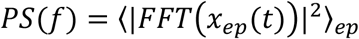

The Normalized Power (NP) at the tag frequency was calculated as the ratio between the power spectrum at the tag frequency and the background power, i.e. the value at the tag frequency of the fit of the power spectrum estimated from the neighboring frequency bins. Two fitting methods were compared: the mean and the power-law fit, computed by fitting a line to the logarithm of the power at the neighboring frequency bins (MATLAB function *Polyfit*).

#### 2.3.2 Evoked Power Spectrum

To compute the EPS, the signal was segmented in non-overlapping epochs of length *wl* time-locked to the onset of a stimulation cycle, and then averaged over all epochs. The evoked power spectrum was then computed as the power spectrum of the average:

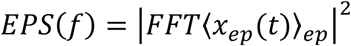

As for the PS, the Normalized Evoked Power (NEP) at the stimulation frequency was calculated as the ratio between the power spectrum at the tag frequency and the background power, i.e. the value at the stimulation frequency of the fit of the power spectrum estimated from the neighboring frequency bins. As for the PS, two fitting methods (mean and power-law) were compared.

#### 2.3.3 Inter-Trial Coherence

As for the EPS, the first step to compute the ITC is to segment the signal in non-overlapping epochs of length *wl* time-locked to the onset of a stimulation cycle. The ITC (also denoted as phase-locking value (Tallon-Baudry et al., 1996)) is computed for each frequency as the vectorial average over all epochs of the phase vectors in complex space, obtained by dividing the Fourier transform of the signal by its amplitude:

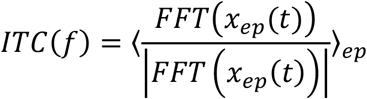

ITC values range between 0 (random phases, corresponding to no coherence) and 1 (perfectly aligned phases, corresponding to maximum coherence). We also tested a normalized version of ITC, where ITC values at the stimulation frequency are normalized by dividing them by the average of the ITC at the nearest frequency bin neighbours.

### 2.4 Performance analysis on simulated data

We assessed the performance of each spectral measure by using the Area Under the Curve (AUC) of the Receiver Operating Characteristic (ROC) curve (Fawcett, 2006), following the same logic used in (Basti et al., 2022) to evaluate the performance of brain connectivity measures with varying data length.

For each SNR value and stimulation frequency, we generated 1000 segments of simulated response, and 1000 segments of simulated rest (see Section 2.2). Each segment was 50 s long. For each set of parameters (spectral measure, data length, *wl*, background fit method) we computed *n*pos = 1000 SSEP estimates from the simulated responses, defined as positives (Fawcett, 2006) and *n*neg = 1000 SSEP estimates from the simulated rest as negatives. We defined a threshold *T*(*p*), which is a function of *p* (0 *< p* ⩽ 100), as the SSEP value at the *p*th percentile of the SSEP distribution estimated from simulated rest. Given a value for *p*, all the scores exceeding *T* (*p*) were classified as positives (detection of SSEP), otherwise as negatives (no SSEP detected); the true positive rate (TPR) was computed as the ratio between the number of positive samples correctly detected as positives and the total number of positives; the false positive rate (FPR) was computed as the ratio between the number of negative samples incorrectly detected as positives and the total number of negatives (in this simulation, FPR is equal to 1−*p*).

We computed the ROC curve by using the Matlab function *perfcurve* (The MathWorks Inc., Natick, Massachusetts) returning the coordinates TPR against FPR for successive values of *p* in the range from 0 to 100 and the associated AUC. We estimated the 95% confidence interval for AUC under large sample size approximation as proposed in (Hanley & McNeil, 1982) (Eq. 1, pag. 32). We assumed the minimum data length required for accurate discrimination as the one for which AUC is larger than 0.70, which is conventionally used as a threshold value to define a diagnostic test as accurate (Swets, 1988).

### 2.4 Statistical performance on real EEG data

To assess the statistical performance of the spectral measures in discriminating the response data from the resting state data for each set of parameters (spectral measure, data length, *wl*, background fit method), we computed the average of the SSEP over a wide posterior cluster of electrodes, *pROI* = (Oz, O1, O2, POz, PO3, PO4, PO7, PO8, Pz, P1, P2, P3, P4, P5, P6, P7, P8), to insure that we captured the low-level visual response to oscillatory on-off patterns (Norcia et al., 2015) within subject-by-subject variability. We then computed a paired Wilcoxon signed rank test between the averaged SSEP during stimulation and during rest across subjects.

## 3 RESULTS

### 3.1 The effect of decreasing data length on SSEP

We start by illustrating the effect of decreasing data length on the estimation of SSEP by applying the three spectral measures on data from one representative subject corresponding to the two stimulations frequencies of the on-off presentation of the checkerboard (4 Hz and 0.8 Hz) and to the rest condition. For both stimulation frequencies, we varied the data length from 16 stimulation cycles (generally already sufficient to obtain robust SSEP) to 4 stimulation cycles (a very short data length).

For 4 Hz stimulation frequency and 16 stimulation cycles (corresponding to 4 s) a clear peak arises at the stimulation frequency in all the three measures (Fig. 1, top panels, blue lines). For decreasing data length, this peak becomes gradually shallower and all the three measures become noisier in the neighbouring frequency bins (Fig. 1, top panels, green and yellow lines); for the shortest data length (4 stimulation cycles, corresponding to 1 s) these effects are extreme: in this example, the peak extends to the neighbor (lower) frequency bin for PS and EPS, and the frequency profile of the ITC becomes flat (Fig. 1, top panels, red lines). At the same time, decreasing data length also has the effect of increasing the instability of the frequency profile of the rest data: while for the longest data length (16 oscillation cycles) the profile around 4 Hz is quite smooth (Fig. 1, bottom panels, blue lines), it becomes noisier for the shortest data lengths (Fig. 1, bottom panels, green, yellow and red lines; note the spurious peak at 3 Hz for PS and EPS). Notably, rest ITC at 4 Hz systematically increases with decreasing data length (Fig. 1, bottom-right panel).

**Figure 1:**
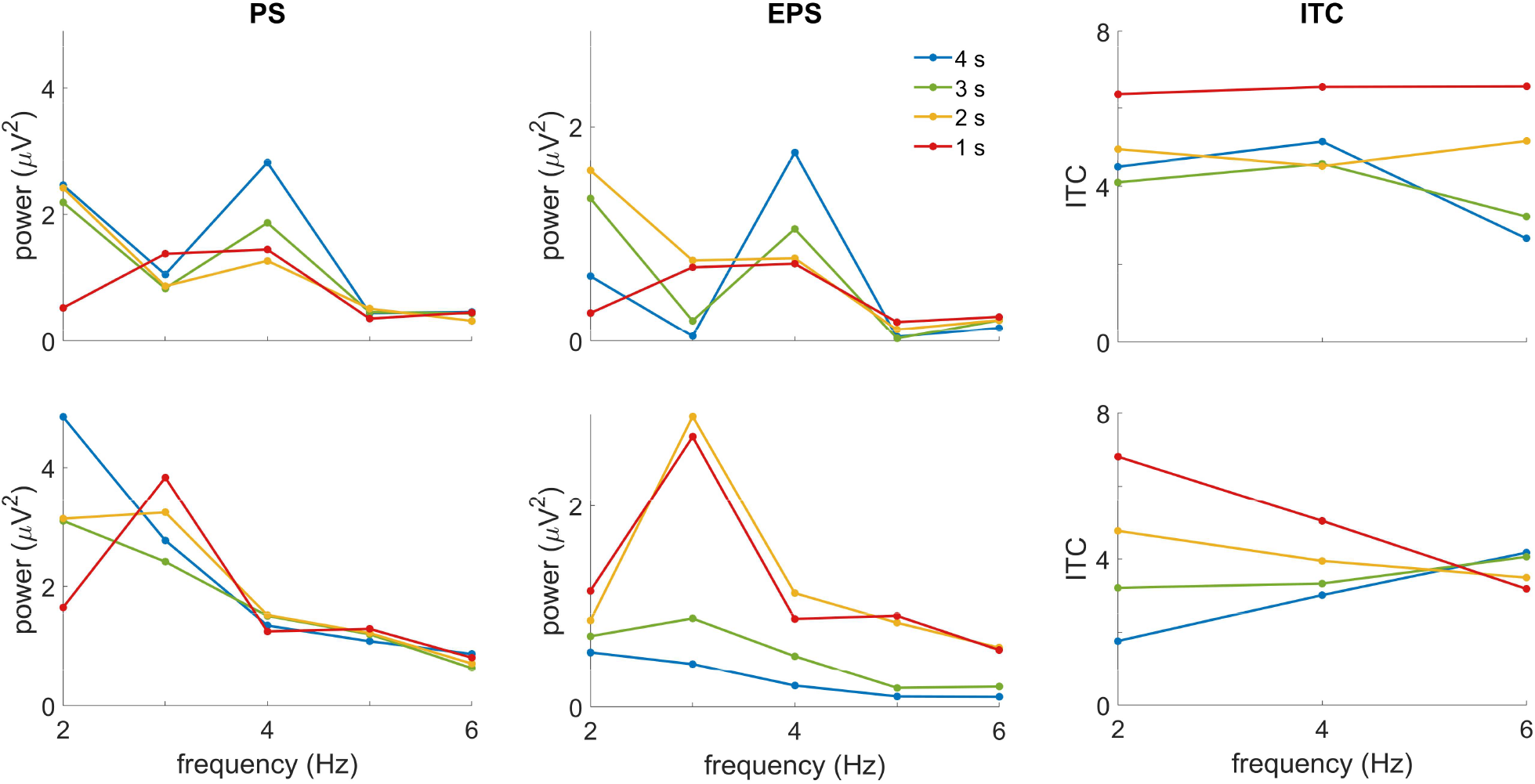
The effect of decreasing data length on SSEP from one representative subject, stimulation frequency = 4 Hz. All figures show the behavior of each spectral measure on data segments decreasing from 4 s (16 cycles of stimulation) to 1 s (4 cycles). Top row: data during stimulation. Bottom row: data during rest. Left-hand column: PS (window length *wl* = 1 s, corresponding to 4 cycles). Middle column: EPS (*wl* = 1 s, corresponding to 4 cycles). Right-hand column: ITC (*wl* = 0.5 s, corresponding to 2 cycles).

Similar effects emerge for 0.8 Hz, with some notable differences compared to 4 Hz: 1) even for the longest data length (16 cycles corresponding to 20 s), PS and EPS peaks are shallower (Fig. 2, left and center top panel); 2) Due to the 1/f spectral profile of the ongoing EEG, PS systematically increases for decreasing frequencies, an effect that often increases with decreasing data length (Fig. 2, left top panel); 3) The ratio between the amplitude of the fluctuations of the spectral measures on rest data and on 0.8 Hz data is higher than for 4 Hz (compare PS and EPS between bottom and top panels, Fig. 2 and Fig. 1, respectively).

**Figure 2:**
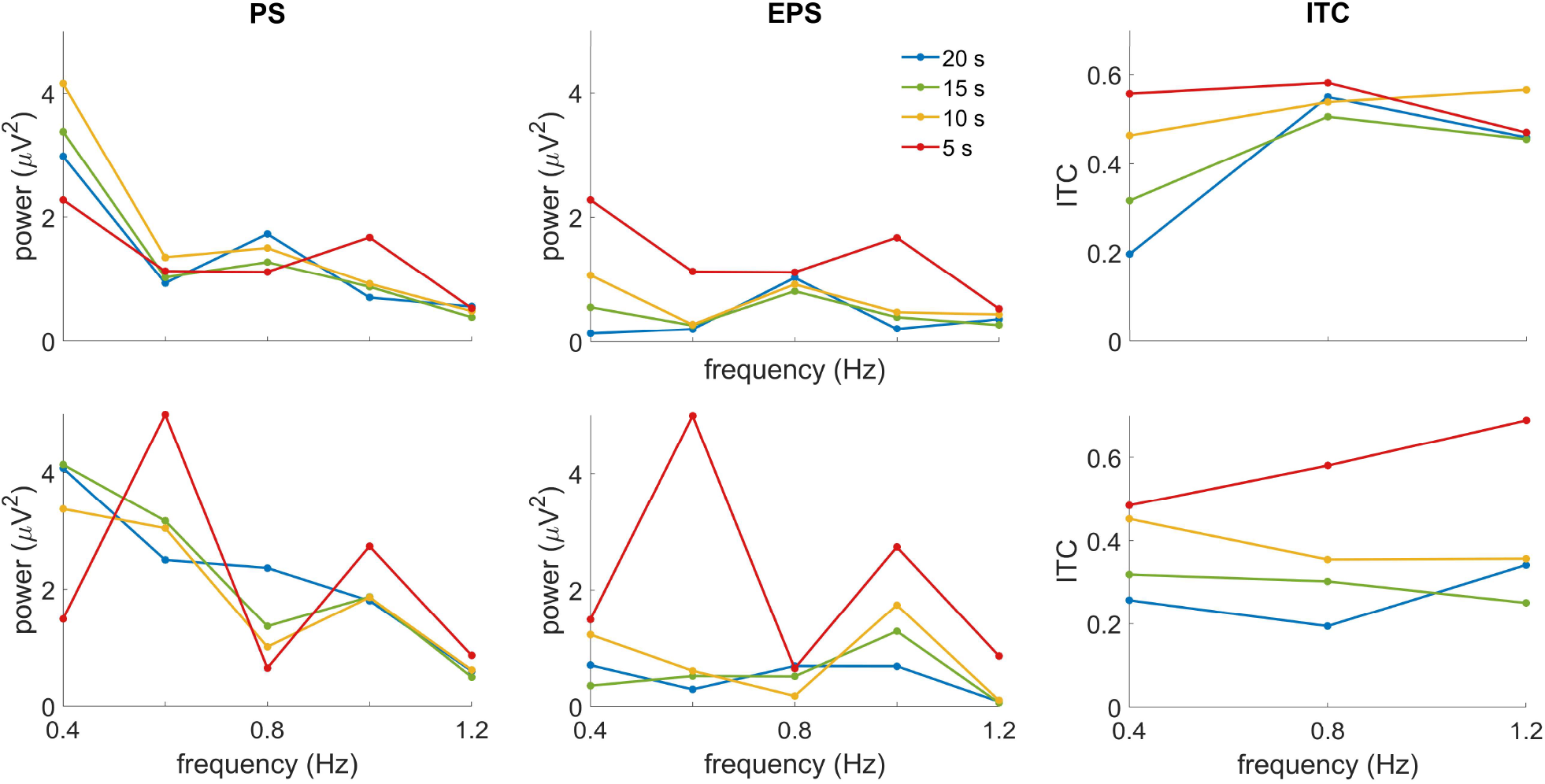
The effect of decreasing data length on SSEP from one representative subject, stimulation frequency = 0.8 Hz. All figures show the behavior of each spectral measure on data segments decreasing from 20 s (16 cycles of stimulation) to 5 s (4 cycles). Top row: data during stimulation. Bottom row: data during rest. Left-hand column: PS (window length *wl* = 5 s, corresponding to 4 cycles). Middle column: EPS (*wl* = 5 s, corresponding to 4 cycles). Right-hand column: ITC (*wl* = 2.5 s, corresponding to 2 cycles).

### 3.2 The influence of spectral measure, window length and spectral fit on SSEP estimation: results from simulated data

We first tackled the question of the optimal measure and parameters to estimate the SSEP in the limit of short data length by using a generative model that simulates the EEG signal as the sum of a noisy stimulus-related response with a stimulus-independent background component. In order to compare the results stemming from the model with the ones measured from the real EEG data, the basic properties of the simulated data matched the ones of the EEG datasets tested in this work: sampling rate, stimulation frequency, event-related potentials and SNR for the SSEP component, low-frequency power-law fit of resting state data for the background component (see Methods for details).

To evaluate the performance of each spectral measure in estimating the SSEP, for each stimulation frequency (4 Hz and 0.8 Hz) and for each of three different values of SNR (see Methods) we generated two classes of data: a “positive” class consisting of 1000 EEG signals simulating the EEG response to a periodic stimulation (generated as described above), and a “negative” class consisting of 1000 EEG signals simulating the EEG during rest, obtained by generating the background component only.

We quantified the classification performance of each measure with the Area Under the Curve (AUC) of the Receiver Operating Characteristics (ROC) curve (Fawcett, 2006) built on the two classes in function of data length.

We first tested, separately for each spectral measure, the influence of the sliding window length *wl* on the classification performance. For 4 Hz stimulation frequency, the performance of the PS systematically increased with the *wl*: the largest *wl* (8 oscillation cycles, corresponding to 2 s) provided an accurate discrimination between stimulation and rest already with a data length of 2 s, i.e. with one single power spectrum measure, and increased steadily with data length up to 0.98 at 10 s (Fig. 3, top left panel). Performance was systematically lower for the other two *wl* values. Notably, PS with the shortest *wl* performed worse than with longer *wl* even for the shortest data length for which PS with the longer *wl* is computed on half zero-padded data (1 s, corresponding to 4 cycles).

**Figure 3:**
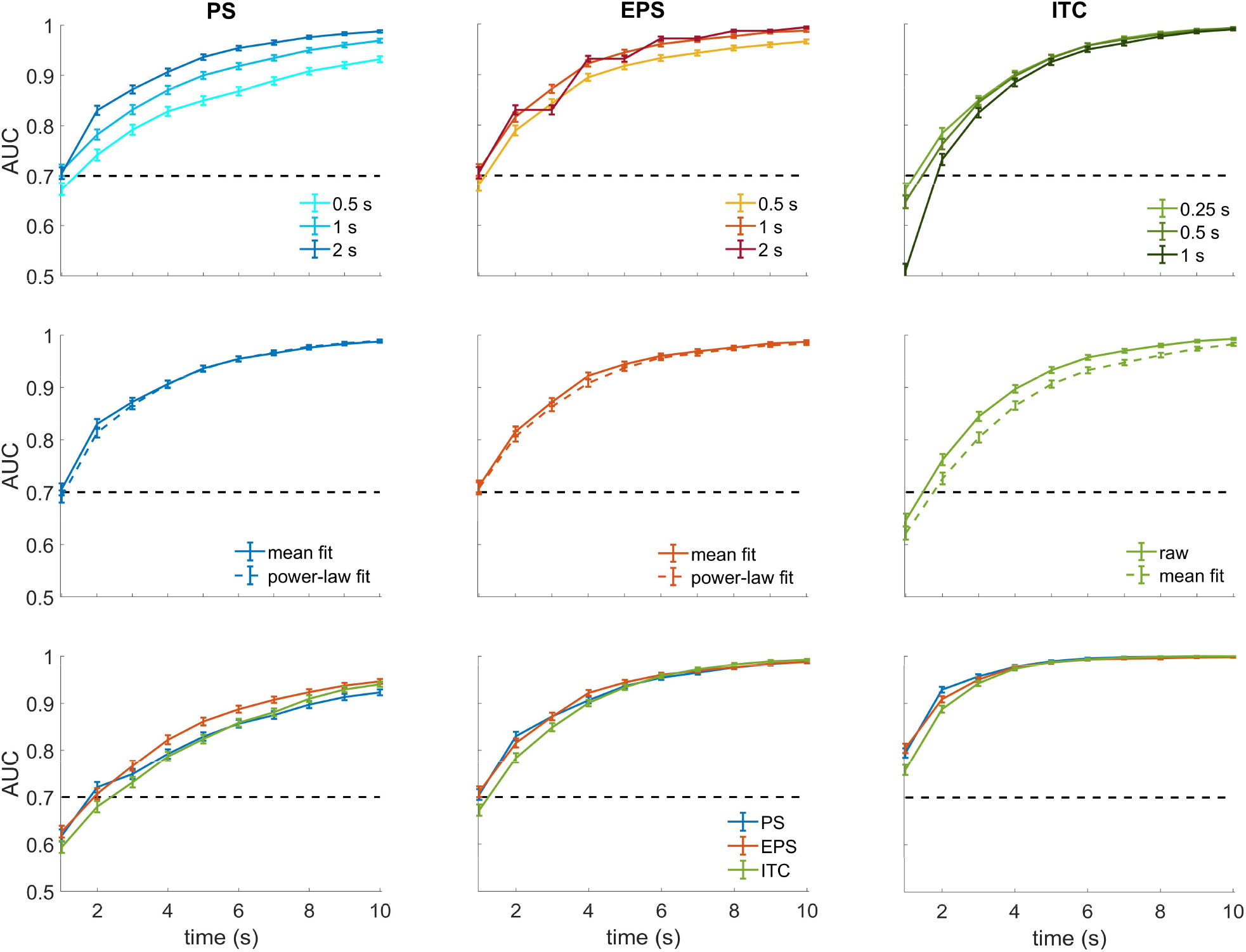
The influence of spectral measure, window length and spectral fit on SSEP estimation from simulated data, stimulation frequency = 4 Hz. All figures show the performance of each spectral measure on data segments decreasing from 10 s (40 cycles of stimulation) to 1 s (4 cycles). Top row: AUC in function of window length *wl*. Middle row: AUC in function of background fit method. The columns of these two rows show the AUC curves for the three spectral measures: PS (left-hand), EPS (middle), ITC (right-hand). Bottom row: AUC of the three spectral measures with their “best” *wl* (2 s for PS, 1 s for EPS, 0.25 s for ITC) in function of SNR. Middle panel: SNR estimated from the real data (the same of the results shown in the top and middle row). Left-hand panel: SNR reduced by 40 %. Right-hand panel: SNR increased by 40 %.

The performance of EPS shows a very similar pattern to PS (Fig. 3, top center panel) but with an important difference: since EPS is computed on non-overlapping windows, for the largest *wl* (8 oscillation cycles, corresponding to 2 s) it improves estimation only at 2 s steps of increase of data length. This limits its performance for short, variable data lengths typical of short recordings as developmental or clinical EEG.

ITC estimation shows an inverse dependence on data length: its performance increases with decreasing *wl*, an effect most probably due to the fact that ITC relies on estimating the phase distribution, which requires as many phase samples as possible (Fig. 3, top right panel).

We then tested the impact of the power-law fit vs the simpler mean fit over the neighbouring frequency bins for estimating the background power in the case of PS and EPS. In both tests, the power-law fit yielded substantially equivalent (though slightly worse) results than the mean fit (Fig. 3, middle left and center panel). Normalizing the ITC by its mean on the closes neighbor frequency bins resulted in a significantly lower performance (Fig. 3, middle right panel).

As a final test, we compared the AUC of each measure with the *wl* associated to the best performance, as a function of SNR (Fig. 3, bottom panel). For high SNR, all the three measures showed a similarly high performance, with slightly worse results of ITC for the shortest data lengths: they all show successful discrimination (AUC > 0.7) already for the shortest length (4 cycles, corresponding to 1 s) and all crossing the value of 0.9 already for length of 12 cycles (3 s). As SNR decreased, the overall performance decreased: The shortest data length allowing discriminatory performance for all the three measures increased to 8 cycles (2 s) for medium SNR and 12 cycles (3 s) for low SNR. Notably, for the lower SNR, ITC showed lower performance for the shortest data lengths, while PS performed systematically worse than the other two measures as data length increases.

For 0.8 Hz stimulation frequency, results were remarkably similar, showing the same dependence on *wl* of the three measures: performance increases with longer *wl* for PS and EPS, while the opposite occurs for ITC (Fig. 4, top row). The power-law fit was equivalent to the mean fit for PS, while slightly but consistently worse for EPS, and the mean fit was clearly detrimental for ITC (Fig. 4, middle row).

**Figure 4:**
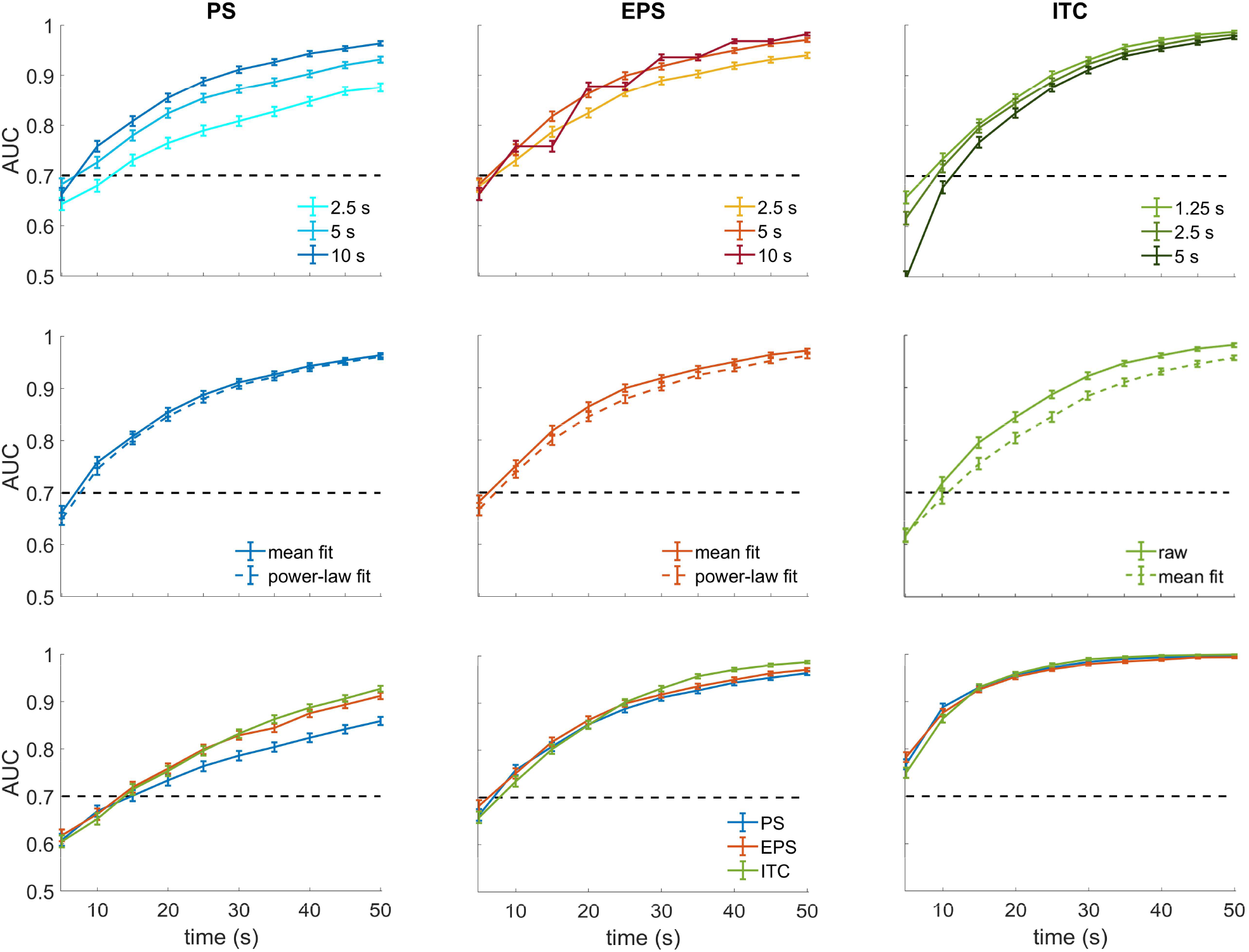
The influence of spectral measure, window length and spectral fit on SSEP estimation from simulated data, stimulation frequency = 0.8 Hz. All figures show the performance of each spectral measure on data segments decreasing from 50 s (40 cycles of stimulation) to 5 s (4 cycles). Top row: AUC in function of window length *wl*. Middle row: AUC in function of background fit method. The columns of these two rows show the AUC curves for the three spectral measures: PS (left-hand), EPS (middle), ITC (right-hand). Bottom row: AUC of the three spectral measures with their “best” *wl* (10 s for PS, 5 s for EPS, 1.25 s for ITC) in function of SNR. Middle panel: SNR estimated from the real data (the same of the results shown in the top and middle row). Left-hand panel: SNR reduced by 40 %. Right-hand panel: SNR increased by 40 %.

All the three measures showed similar results with varying SNR: the minimum data length for successful discrimination (AUC > 0.7) was 4 cycles (5 s) for the higher SNR, 8 cycles (10 s) for the middle SNR and 12 cycles (15 s) for the lower SNR (Fig. 4, bottom row). The most notable difference with 4 Hz is that for the lower SNR the difference between PS and the other two measures is more important, starting already from 16 cycles (20 s).

### 3.3 The influence of spectral measure, window length and spectral fit on SSEP estimation: results from real data

We then tested whether the results obtained with the simulated data reflected the behavior of the spectral measures on real data. We measured the performance of the three spectral measures in discriminating between data segments during periodic stimulation and rest segments by computing the statistical significance (paired Wilcoxon test) of the difference between stimulation and rest condition across all subjects (see Methods).

The analysis on 4 Hz stimulation data shows results mostly similar to simulated data: all the three measures show a significant discrimination already for data length of 8 cycles (2 s) (Fig. 5, top row), but while ITC for the shortest window lengths shows the same pattern (short *wl* yield higher statistical significance than longer ones), the dependence on *wl* of PS and EPS is more heterogeneous. As for the simulated data, the alternative power-law spectral fit showed almost equivalent performance for PS and EPS, and the mean spectral fit proved detrimental for ITC (Fig. 5, bottom row). 0.8 Hz real data behaved very closely to simulated data: the dependence on *wl* followed the same pattern (higher performance of longer windows for PS and EPS, the opposite for ITC, Fig. 6, top row); the power-law spectral fit yielded almost identical scores for PS and EPS, while the mean fit for ITC was detrimental (Fig. 6, bottom row); statistical significance was reached for all the three measures already for data length of 8 cycles (10 s).

**Figure 5:**
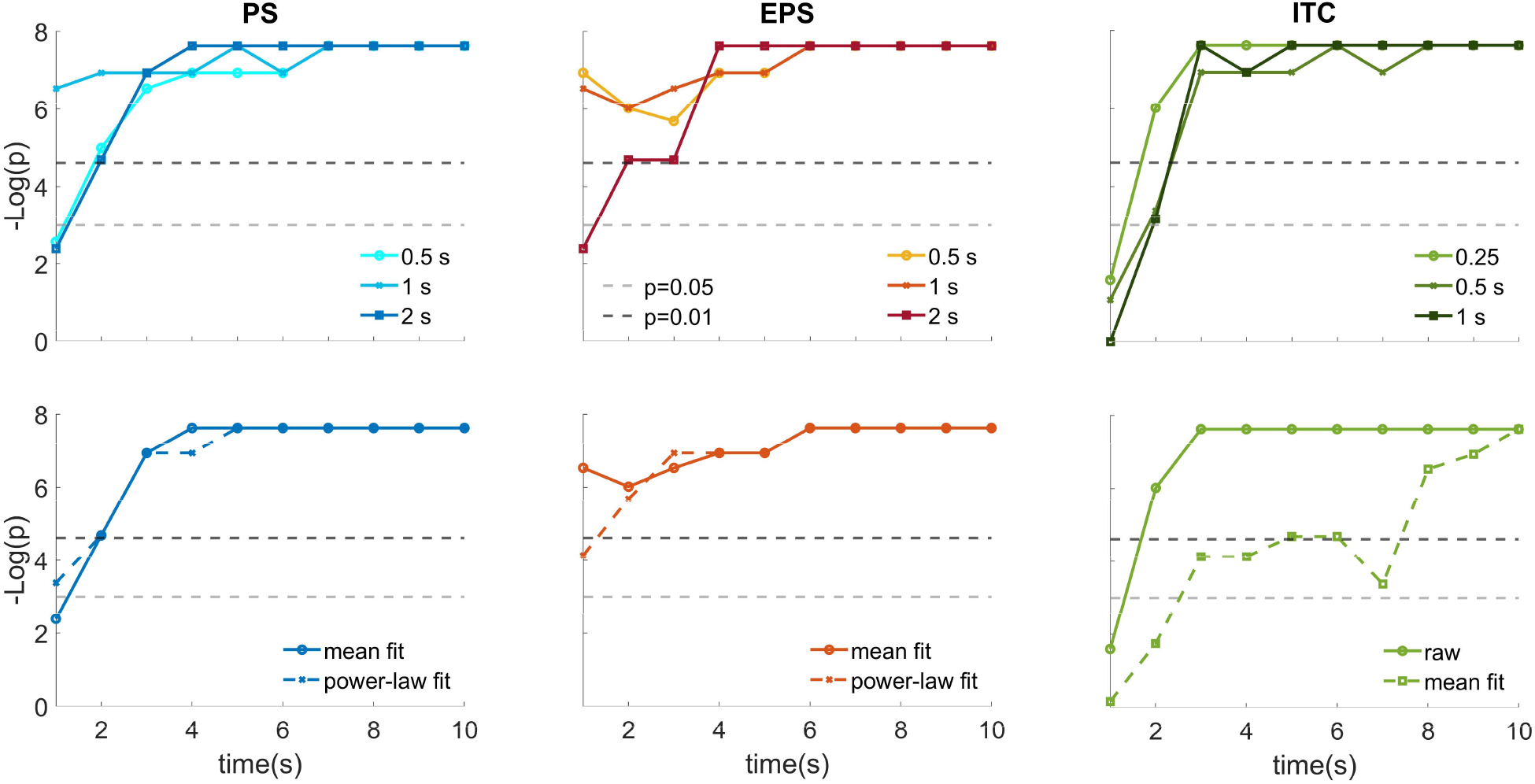
The influence of spectral measure, window length and spectral fit on SSEP estimation from real data, stimulation frequency = 4 Hz. All figures show the statistical performance of each spectral measure in discriminating between stimulation and rest, quantified with -log(*p*) (where *p* is the p-value of the Wilcoxon paired rank test between stimulation and rest across subjects) on data segments decreasing from 10 s (40 cycles of stimulation) to 1 s (4 cycles). Top row: Statistical performance in function of window length *wl*. Bottom row: Statistical performance in function of background fit method. The columns show the statistical performance for the three spectral measures: PS (left-hand), EPS (middle), ITC (right-hand).

**Figure 6:**
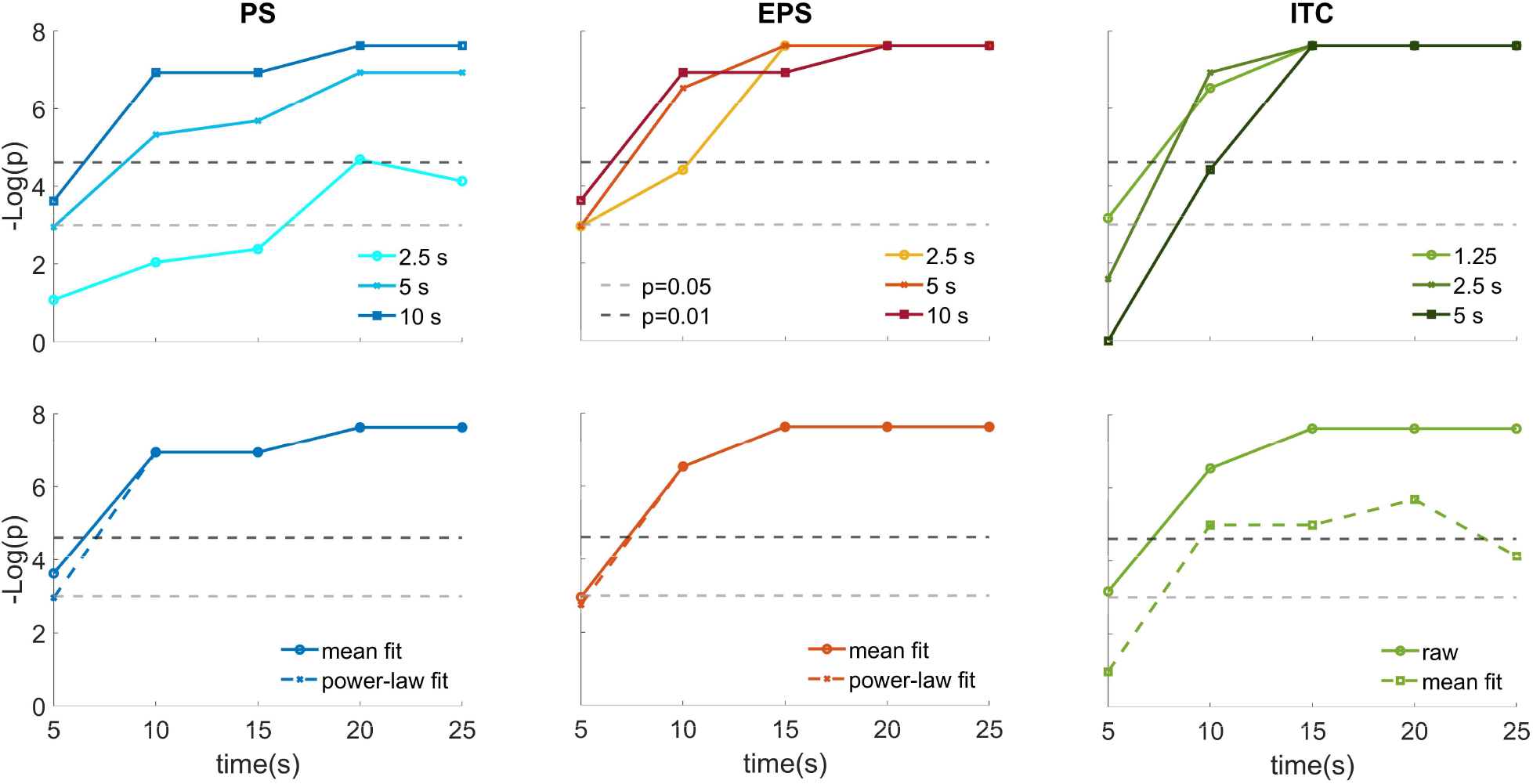
The influence of spectral measure, window length and spectral fit on SSEP estimation from real data, stimulation frequency = 0.8 Hz. All figures show the statistical performance of each spectral measure in discriminating between stimulation and rest, quantified with -log(*p*) (where *p* is the p-value of the Wilcoxon paired rank test between stimulation and rest across subjects) on data segments decreasing from 25 s (20 cycles of stimulation) to 5 s (4 cycles). Top row: Statistical performance in function of window length *wl*. Bottom row: Statistical performance in function of background fit method. The columns show the statistical performance for the three spectral measures: PS (left-hand), EPS (middle), ITC (right-hand).

A direct comparison of the results of the two stimulation frequencies using the “best” *wl* for each spectral measure shows that in both conditions the three measures show equivalent performance that depends on the number of cycles of the available data: the minimum length for obtaining a reliable measure (*p*<0.01) is 8 cycles (Fig. 7).

**Figure 7:**
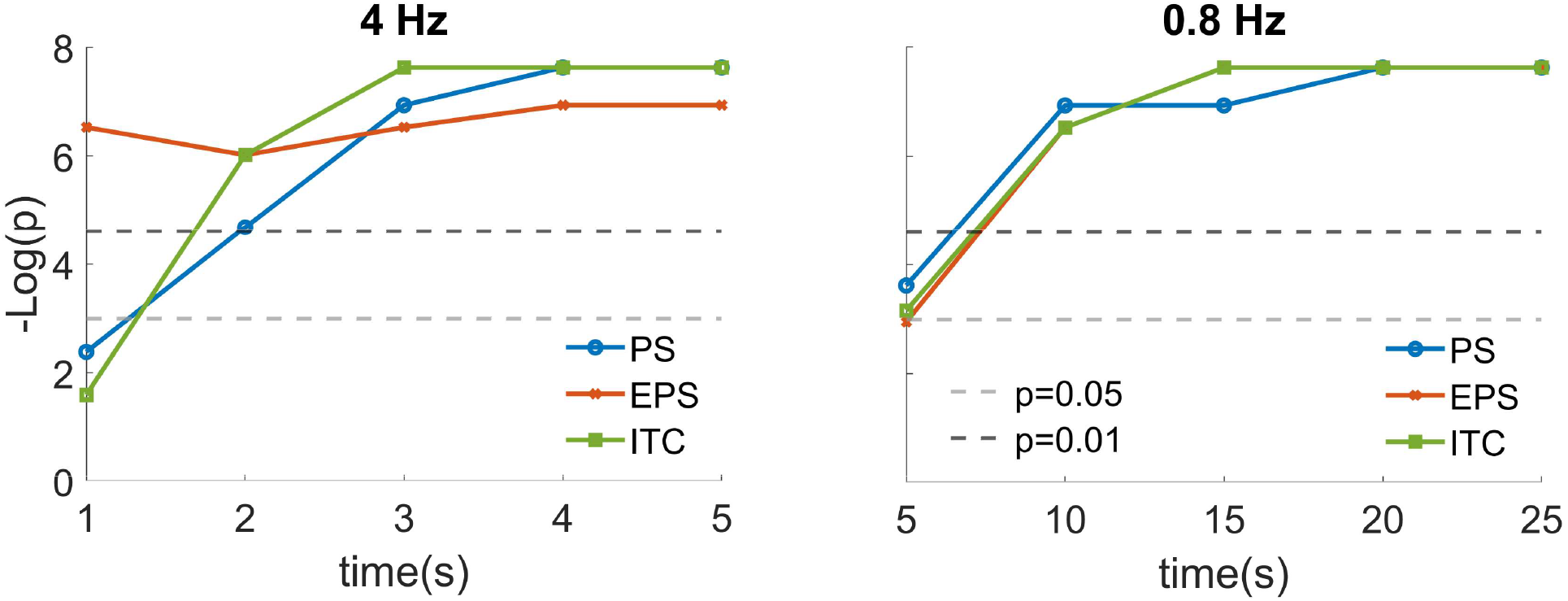
SSEP estimation from real data in function of stimulation frequency. Statistical performance of the three spectral measures with their “best” *wl* (8 cycles for PS, 4 cycles for EPS, 1 cycle for ITC) for stimulation frequency of 4 Hz (left-hand panel) and 0.8 Hz (right-hand panel).

## 4 DISCUSSION

We investigated with both data simulations and SSVEP measured from a real EEG dataset the methods of analysis and the parameters that most efficiently compute SSEP at low stimulation frequencies in the limit of short data. This analysis also provided an estimate of the minimum data length necessary to obtain a reliable response for a given stimulation frequency. Here we report the main results of our analysis:

- Performance of all the three spectral measures crucially depended on *wl*, the length of the sliding time window used to compute each measure: for PS and EPS, longer *wl* provided better performance, while the opposite occurred for ITC. This effect was more relevant for shorter window lengths and for PS than for the other two measures.
- Once the optimal *wl* was set, at short data length the three measures showed equivalent performances.
- For increasing data length and low SNR, PS performance increased significantly slower than EPS and ITC, while the latter two measures were substantially equivalent;
- For the SNR levels corresponding to the real SSVEP analyzed in this paper, the minimum data length necessary to obtain a reliable response was 8 cycles of stimulation frequency, both for 0.8 Hz and 4 Hz.
- The analyses on the real data confirmed the main results emerging from the simulation, in particular for the 0.8 Hz stimulation frequency, suggesting that the SSEP generative model we implemented is valuable and may be useful for further investigations.

Hereafter we comment on the potential causes underlying these results, on their impact on studies involving short recordings and on their relation with respect to the current state-of-the-art of SSEP estimation.

### Which measure in the limit of short data length?

The dependence of ITC on *wl* for short data length is most likely due to the finite-size effect of measuring ITC: fewer phase samples cause the phase distribution to be heterogeneous even for rest data, eliciting a spurious increase of ITC (Fig. 1 and 2, bottom right panel). This dependence on *wl* rapidly fades away for increasing data length, where the number of samples is sufficient to draw a homogeneous null distribution.

The dependence of PS and EPS on *wl* for short data length is less obvious: it suggests that frequency resolution is more relevant than averaging, even when this means computing the power spectrum on a single time window.

In practice, these results show that in the limit of short data length the three measures are equivalent, and may provide a valuable result even for very short data segments (8 cycles, corresponding to 10 seconds for the lowest frequency, 0.8 Hz), provided that the largest possible *wl* is used in the case of PS and EPS, and the opposite for ITC. Therefore, if the stimulation elicits a significantly high SNR, a few tenths of seconds are sufficient to measure reliable SSEP. This observation is particularly important for developmental studies, where the attentional span of the subjects is very limited. For example, they support the reliability of the analyses implemented in (Buiatti et al., 2019), where we presented visual stimuli exactly at a frequency of 0.8 Hz and we analyzed the SSEP by computing PS on data as short as 15-20 s per condition for some subjects. For this kind of studies where the segments to be analyzed are short and variable, PS adapts more efficiently than EPS and ITC to their variable length for two reasons: 1) while epochs for the computation of EPS and ITC need to start at the onset of a stimulation cycle, this is not necessary for the PS because it is independent of the phase; 2) differently from EPS and ITC, the PS may be computed with overlapping windows.

An alternative method of analysis that might be successful in these conditions is Canonical Correlation Analysis (CCA, (Lin et al., 2006)). We have recently shown that a normalized version of CCA is very efficient (and better than PS) in detecting SSVEP from low-density recordings with low-frequency stimulations (Kartsch et al., 2022). Further research is welcome to test the performance of CCA on SSEP with low-frequency stimulations.

### Which measure for long data lengths?

While this work focused on SSEP estimation from short data length, our simulations suggest that for long data length (or, equivalently, many trials), EPS and ITC perform increasingly better than PS, particularly for low SNR (Figs. 3 and 4, bottom right panel). This result supports the much wider use of EPS and ITC than PS for estimating SSEP in studies where data statistics is not an issue (e.g. (Ding et al., 2016; Henin et al., 2021; Rossion et al., 2015)). This result might also explain the ones emerging from the simulation of low frequency (1.1 Hz and 3.3 Hz) SSEP performed by (Benjamin et al., 2021), where PS has a drastically lower performance than EPS and ITC: their simulation consists in much longer data with a much lower SNR than those used in our work, and a simulated fixed-latency response that favours measures of phase-locked activity.

In general, SNR largely varies between different studies focusing on different brain responses. The results presented in this work are based on a visual stimulation (on-off checkerboard) that elicits a high SNR. It is likely that for designs eliciting lower SNR, longer data lengths are needed to obtain reliable results, thus supporting the use of EPS and ITC. Establishing an independent measure of SNR on each MEEG dataset would help researchers to choose the optimal measure to analyze it.

### Modeling SSEP

We modeled the shapes and latencies of the simulated transient evoked responses by mimicking the ERPs obtained from our EEG dataset. To rule out that the results of the simulations depended on these specific shapes and latencies, we performed the AUC analysis for a variety of other responses, obtaining very similar results.

We modeled SSEP as transient evoked responses because our aim was to explore SSEP estimation in the low frequency limit, where this generative model is generally accepted (Norcia et al., 2015). However, an alternative model proposes that SSEP arise from the phase-locking of ongoing brain oscillations (Makeig et al., 2002). It is possible that the two mechanisms of SSEP generation coexist with variable proportion depending on the relation between the stimulation frequency and intrinsic neural time scales, as well as on the brain activity (driven by sensory input vs by top-down modulations). Testing the results stemming from a model including a phase-locking mechanism would shed more light on these mechanisms. This alternative model might explain the heterogeneity of the dependence on *wl* for 4 Hz SSEP (Fig. 5).

### Spectral fit of the background

Results both from the simulations and from real data show that the power-law fit is equivalent to the mean fit for PS, while it is slightly detrimental for EPS. This result may be explained by the fact that at low frequencies the spectral profile of the PS heavily influenced by the 1/f increase of the ongoing neural activity, therefore in this case the power-law fit is effective. On the other hand, the averaging involved in the computation of the EPS removes a large part of ongoing fluctuations, therefore the power-law is not a good model of the residual background spectrum anymore.

Since the mean fit of the PS provides overestimates the background power (because the power spectrum is much steeper for lower-frequency compared with higher-frequency bins around the stimulation frequency (Buiatti et al., 2019)), our suggestion is to use the power-law fit for the PS and the mean fit for the EPS.

### Methodological indications for the analysis of SSEP with low stimulation frequency

In the light of these results, hereafter we provide some methodological suggestions for the analysis of SSEP with low stimulation frequency (< 6 Hz).

- In general, unless there is a strong prior assumption on the model underlying the analysed SSEP, it is a good practice to test all the three spectral measures (PS, EPS and ITC) as they capture different aspects of the SSEP.
- For long data (> 20 cycles of stimulation), we advise to use EPS and/or ITC, as their performance usually overcomes the one of PS, especially for low SNR.
- For short data (< 20 cycles of stimulation), we advise to use PS and EPS with the largest possible *wl*, while the opposite for ITC. In general, for this data length the three measures are equivalent.
- For short data (< 20 cycles of stimulation) and high SNR, in cases where data length is variable we suggest to use PS, as it adapts better than EPS and ITC to this variability.
- In this low frequency range, we suggest to normalize PS at the stimulation frequency by the power-law fit of PS at neighbouring frequency bins because this normalization provides an unbiased estimate of the peak amplitude; on the other hand, we suggest to use a mean fit for EPS, and no spectral fit for ITC.

## 5 CONCLUSIONS

Our work aims to set a methodological reference for estimating SSEP with low frequency stimulation in the delta frequency range and in the limit of short data length. We provide precise indications on the use of the three main spectral measures of SSEP – power spectrum, evoked power spectrum and inter-trial coherence – in function of stimulation frequency, data length and SNR. While the real data used in this work are EEG recordings, these indications are valid for MEG data too, as the origin of the temporal structure of the two signal is the same.

The overall result is that, for relatively high SNR, with an accurate choice of the parameters it is possible to obtain reliable measures of SSEP to low frequency stimulation even with very short data. This result encourages researchers to use low-frequency stimulation SSEP designs with challenging populations like infants, patients or in out-of-the-lab settings. Given the consistency between the results from simulated and real data, we also emphasize the utility of our SSEP generative model to investigate and calibrate the optimal methods of measure of SSEP (and potentially other kinds of neural signals) on real data.

## Acknowledgements

We thank Giorgio Vallortigara for his generous support to this project.

## Author contributions

MB: conceptualisation; methodology; software; investigation; formal analysis; writing - original draft preparation and review and editing; visualization; supervision. DS: software; investigation; formal analysis.

